# Ancient brainstem inhibitory neurons control selective spatial attention

**DOI:** 10.1101/2022.06.27.497794

**Authors:** Ninad B. Kothari, Arunima Banerjee, Qingcheng Zhang, Wen-Kai You, Shreesh P. Mysore

## Abstract

To behave adaptively in complex environments, animals must selectively process the most important information in space while ignoring distractors. Contrary to the dominant cortico-centric view, we report that an evolutionarily old group of inhibitory neurons in the brainstem, called PLTi, is critical for this function of selective spatial attention. In freely behaving mice performing a human-like spatial attention task, we found that bilateral silencing of PLTi severely disrupted target selection without causing perceptual or motor impairments. PLTi’s effects depended necessarily on goal-relevant, rather than just physical salience-based signals, together revealing it as a dedicated brain site for priority-driven attentional target selection. PLTi’s core computational function is its explicit implementation of the abstract cognitive decision boundary separating target from distracters: it controlled the accuracy and categorical precision of this decision boundary, doing so by shaping neural representations of competing stimuli in the superior colliculus, a major sensorimotor hub. PLTi may, therefore, be a conserved brainstem engine across vertebrates for winner-take-all-like spatial decisions in goal-driven behavior.

## Introduction

Sensory environments are replete with competing sources of information that can guide behavior. The informativeness of each stimulus in the environment for animal behavior arises both from its physical distinctiveness (or salience) - a ‘bottom-up’, exogenously driven component, and its relevance to the goals of the animal (or behavioral relevance) - a ‘top-down’, endogenously driven component^1^. The combined salience and relevance of each stimulus, defined as its ‘priority’, is considered as its net informativeness for guiding behavior^1^. Animals possess the remarkable ability to select and preferentially process the highest priority stimulus in space while ignoring distracting stimuli of lower priority. This ability, termed selective spatial attention, is crucial for adaptive behavior and even survival^1-3^, and is an executive function that is disrupted in a range of neuroatypical conditions including ADHD and schizophrenia^4,5^.

A rich literature of studies in human and non-human primates from over half a century has yielded important insights into the consequences of selective spatial attention to behavior^2^ as well as to neural representations^6,7^. However, how the brain selects the attentional target, in the first place, is unknown. Moreover, in the natural world, the priorities of competing stimuli can vary over a large range owing to their diverse physical and goal-relevant properties, and can vary dynamically from one moment to the next owing to changes in these properties as well as changes in the very composition of stimuli in the environment. As a result, neural circuits underlying selective spatial attention must, necessarily, be able to reliably select the highest priority target not only when it may be easily discernable from lower priority distracters (i.e., when the relative priority between target and distracters is very large), but also when this discrimination may be challenging (i.e., when distracters are close in priority to the target). A fundamental question, then, that has remained wide-open is, “How does the brain select the attentional target among distracters over the natural range of relative priorities?”

The literature has also highlighted key brain regions involved in mediating selective spatial attention^6,8-12^. The dominant view from this work is that selective spatial attention is controlled principally by cortical, fronto-parietal brain networks^6,9^. This view, however, is at odds with emerging evidence of powerful, spatially selective target processing for goal-driven behavior^13^ in vertebrate species with less well-developed fronto-parietal cortices, such as rodents^14-17^, or even absent fronto-parietal cortices, such as birds^18^ and fish^19^. These findings lead to the puzzling neuroethological question, “How, then, might selective spatial attention be implemented in these vertebrate species with poorly developed fronto-parietal networks?”^13^. A distinct possibility is the involvement of an evolutionarily conserved (subcortical) neural substrate^13,20^. Whereas work in primates has identified the midbrain superior colliculus (SC) as being involved in attentional control^21^, whether it is itself the site at which target selection is actually computed^22^, and how, mechanistically, it implements the competitive interactions among stimulus representations that are crucial for its role in spatial attention^21^ are unknown^8,10,20,23^. Moreover, because the SC is a sensorimotor area, as are the key fronto-parietal brain areas identified above, their disruption impairs not only attention control but also a range of co-occurring functions including perception of single stimuli^24-26^, execution of orienting movements^26-28^, and selection among movement plans^29-31^. Consequently, an intriguing open question is, “Are there mechanistic loci in the brain dedicated to implementing competitive target selection and distracter suppression for selective spatial attention?”.

Here, we address these questions and discover a common link that connects them, namely, an evolutionarily old (Supplementary Figure 1)^20,32-34^ group of brainstem inhibitory neurons called PLTi. In freely behaving mice performing a human-like task of selective spatial attention, we demonstrate that PLTi is a powerful, dedicated brain locus for the implementation of priority-dependent distracter suppression and target selection, we uncover PLTi’s computational contributions to attention control, and reveal its mechanistic path of action.

### PLTi is a conserved group of PV+ inhibitory neurons in the mouse tegmentum that functionally inhibits the SC

What might be a potentially conserved subcortical site involved in selective spatial processing for guiding behavior? Converging evidence from work across vertebrate species led us to a group of inhibitory neurons in the brainstem. First, work in behaving monkeys has highlighted a critical role for the superior colliculus, a midbrain sensorimotor hub^10,21,30,35^, in the selection of a spatial target among distracters. Specifically, it is competitive interactions among representations of the target and distractors in the space map of the intermediate and deep layers of SC (SCid) that are necessary for spatial target selection^10,21,30^. The mechanistic source of these competitive interactions underlying behavior, however, is unknown. Second, work in awake, but passive birds (owls and pigeons) has shown that a group of parvalbumin-positive (PV+) inhibitory neurons in the brainstem tegmentum, called Imc (nucleus isthmi pars magnocellularis), drive competitive stimulus interactions across the space map in the avian homolog of the SCid (OTid)^36-38^ (and in sleeping reptiles, competition among brain hemispheres^39^). However, what role they may play in controlling goal-driven behavior is unknown in mammals (and indeed, in any vertebrate species). These inhibitory neurons are considered to lie in the parabigemino-tegmental complex in mammals^32-34^ (Figure 1a, left and middle).

**Figure 1.**
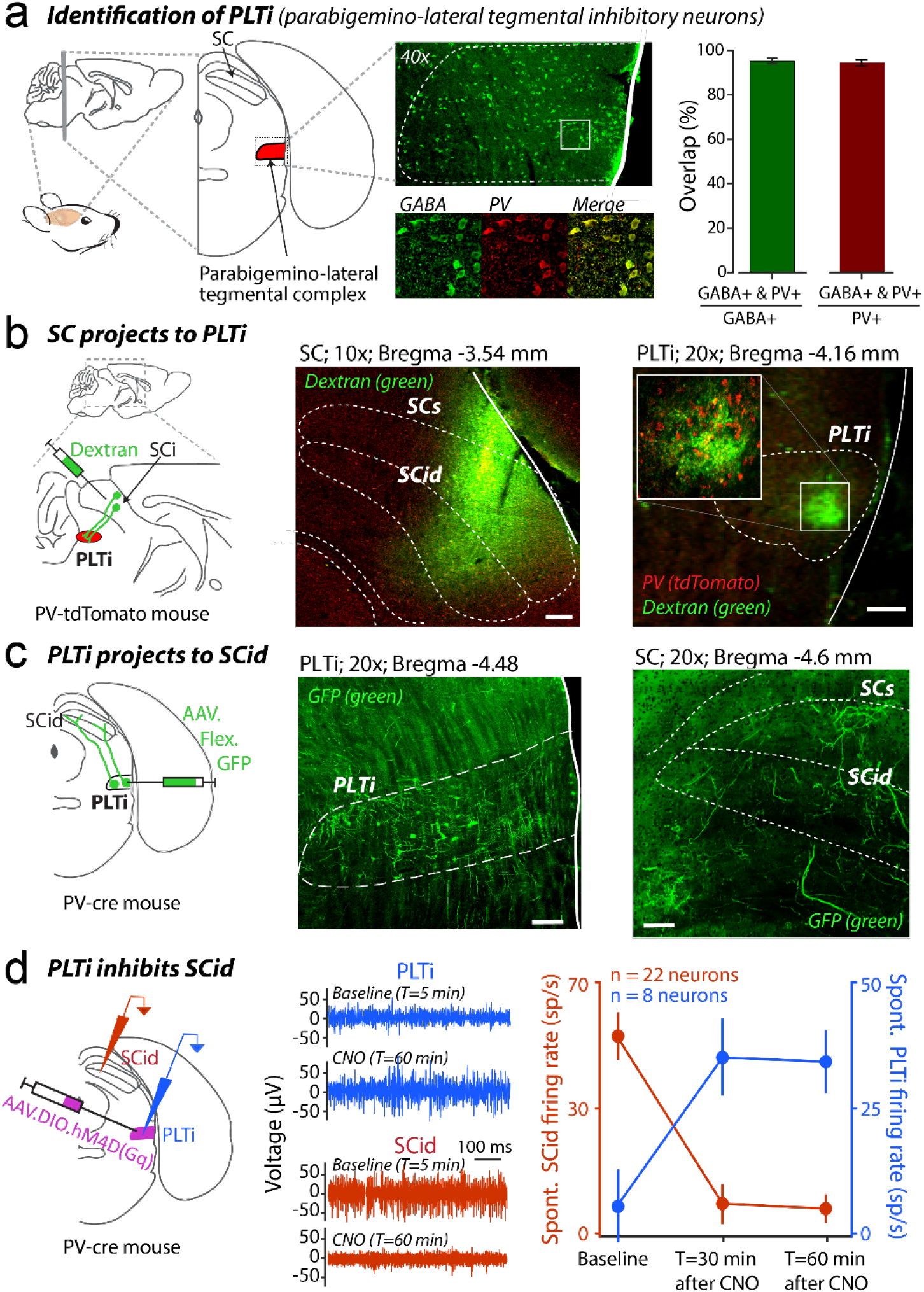
PLTi is a conserved group of PV+ inhibitory neurons in the midbrain parabigemino-lateral tegmental complex that functionally inhibits the SC. **(a)** Identification of PLTi (parabigemino-lateral tegmental inhibitory neurons). (Left) Anatomical location of parabigemino-lateral tegmental complex area (Methods). (Center top) Immuno-histochemical labeling of GABA+ (green) cells in this complex. Dotted line shows the outline of the complex. (Center bottom) Zoomed-in image of colocalized GABA+ (green) and PV+ (red) cells in the complex (arrows indicated cells). (Right) Summary of percentage of GABA+ cells in the complex that were also labeled for PV+ (green bar), and vice-versa (red bar; n=3 mice; see Supplementary Figure 1). (b) SC projects to PV+ neurons in the complex. (Left) Schematic of Dextran injection (green) in SCi (Center) injection site in the SCi (dotted white lines outline SC). (Right) Dense projection fibers (green) in the complex (dotted white lines). (Inset) Zoomed-in view of white square showing SC projection fibers (green) surrounding PV+ cells (red) in the complex. (c) PV+ neurons in the complex project to SCid. (Left) Schematic of location of AAV.Flex.GFP injection (green) in the complex in PV-Cre mouse and projection fibers in the SCid. (Center) Injection site in complex (dotted white lines). (Right) Projection fibers (green) in SCid (white dotted lines). (d) PV+ neurons in the complex functionally inhibit SCid. (Left) Schematic of location of excitatory chemogenetic injection (pink: AAV.DIO.hM3D(Gq)) in PV-Cre mouse and locations of simultaneous recordings of extracellular activity at injection site (blue) and SCid (red). (Center) Bandpass filtered extracellular activity of paired recordings from PV+ neurons in the complex (top, blue) and SCid (bottom, red). (Right) Mean activity of isolated units (n=22, two mice) in SCid measured in the baseline condition and at time points after ip injection of CNO (PLTi activated condition). All scale bars are 100 μm unless indicated. See also Supplementary Figure 1.

To identify these inhibitory neurons in the mouse parabigemino-tegmental complex, we immunostained for GABA in midbrain sections. We found GABA+ cells distributed across this region (Fig.1A, middle; Methods), with nearly all GABA+ cells in the parabigemino-tegmental complex also labeled for parvalbumin (PV) (Figure 1a right – red bar plot, 95.01% ± 0.46% GABA+ cells were also PV+; Figure 1a right – green bar plot, 94.33% ± 1.33% PV+ cells were also GABA+). Injection of a fluorescent tracer (Dextran; Methods) focally into the SC (Figure 1b left, middle) showed anterogradely labeled fibers focally within the parabigemino-tegmental complex (Figure 1b-right). Conversely, injection of a cre-dependent AAV vector expressing eGFP selectively in the PV+ cells of the parabigemino-tegmental complex (60 nL; AP-4.30; ML+1.75; DV-3.5; Figure 1c, left and middle) showed anterogradely labeled fibers broadly across the intermediate and deep layers of the SC (Figure 1c, right). Finally, we selectively activated these PV+ neurons in awake, but passive (head-fixed) mice using chemogenetics (Figure 1d left; administration of chemogenetic ligand CNO vs. saline to mice expressing the excitatory chemogenetic receptor, hM3Dq, stereotactically in PV+ neurons; Methods). This produced suppression of spontaneous activity in SCid, revealing that these PV+ neurons functionally inhibit SCid (Figure 1d middle and right; red, neurons in SCid: baseline: 47.01 +/- 11.31 sp/s; 30 mins after CNO: 7.2 +/- 9.4 sp/s; 60 mins after CNO: 5.9 +/- 6.82 sp/s; blue, PV+ neurons in parabigemino-tegmental complex: 5.39 +/- 14.57 sp/s; 30 mins after CNO: 35.05 +/- 15.24 sp/s; 60 mins after CNO: 34.2 +/- 12.56 sp/s Methods). Thus, these PV+ neurons in the parabigemino-tegmental complex receive inputs from SC, send long-range projections to SCid, and suppress SCid activity (analogously to the avian Imc neurons^20,37,40^). We refer to this group of PV+ GABAergic neurons in the mouse as *parabigemino-lateral tegmental inhibitory neurons* (**PLTi,** ^32,33^).

### PLTi controls priority-driven spatial target selection

Does the PLTi play a role in mediating selective spatial attention? To test this, we first trained freely behaving mice on a modified version of our recently developed touchscreen-based flanker task^14^ (Figure 2a, Supplementary Figure 2a). This task in mice is modeled directly after the classic flanker task of selective spatial attention in humans, which is known to operate on stimulus priorities, i.e., on the combination of the stimulus-driven physical salience and goal-driven behavioral relevance of each stimulus^41,42^. Briefly, we trained mice to report the orientation (vertical or horizontal) of a central target grating, while simultaneously ignoring the orientation (also vertical or horizontal) of a flanker grating, located peripherally (Figure 2a; Methods). Mice were rewarded for correctly reporting the orientation of the target with a nose-touch at one of two response ports located along the elevational axis (Figure 2a; horizontally (vertically) oriented target ◊ touch at lower (upper) response port; counterbalanced across animals). This task design allowed the dissociation of the loci at which task-relevant information was present (central or peripheral azimuthal locations) from the loci of behavioral reports (upper or lower response ports). Three task conditions were interleaved randomly: (a) *Target-alone condition*, in which the flanker was absent, (b) *Congruent flanker condition*, in which the target and flanker had the same orientation, and (c) *Incongruent flanker condition*, in which the target and flanker had opposite orientations (Figure 2a, Supplementary Movie 1). The contrast of the target was always fixed while that of the flanker (when present) was varied parametrically from being lower to higher than that of the target, allowing us to systematically vary relative priority between the target and distractor (Methods^14^).

**Figure 2.**
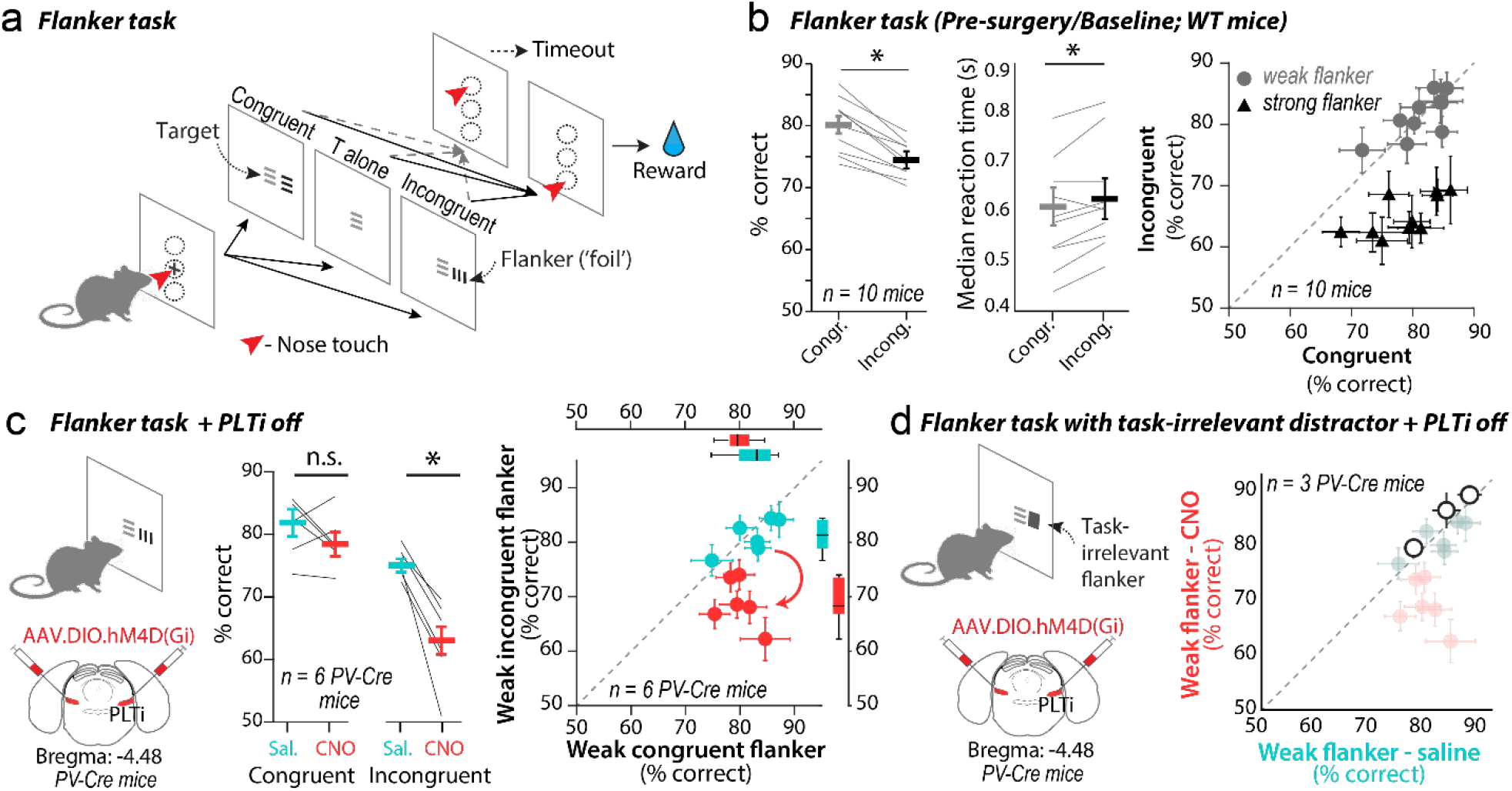
PLTi controls priority-dependent spatial target selection. **(a)** Schematic of flanker task. Vertically arranged black circles in left and right most panels show nose-touch mask used in the behavioral apparatus (See also Supplementary Figure 2a). Solid black arrows: choices leading to water reward; horizontal (vertical) target ◊ touch at lower (upper) response port, counterbalanced across mice. Dashed gray lines: choices leading to timeouts. **(b)** Comparison of target selection performance across the congruent and incongruent task conditions pooled across contrasts; n= 10 mice. (Left) Mean percentage correct (p=4.85*10^-5^, t(9)=3.87, paired two-tailed t-test with Holm-Bonferroni correction (HBMC) for multiple comparisons). (Center) Median RT (p=0.019, t(9)=2.43, paired two-tailed t-test with HBMC); each line represents data from one mouse. (Right) Percentage correct (mean +/- s.e.m) for the weak flanker (gray circles) and strong flanker (black circles), congruent and incongruent conditions (2-way RM-ANOVA, p=2.38*10^-13^ for Weak vs Strong flanker followed by HBMC between Weak and Strong for % correct: Congruent - p=0.268, t(18)=1.156; Incongruent - p=3.54*10^-9^, t(18) = 10.61, followed by paired two-tailed t-tests; (see Supplementary Table 1); each point - one mouse. **(c)** (Left) Abbreviated schematic of flanker task, and schematic of bilateral injection of inhibitory chemogenetic virus in PLTi of PV-Cre mouse (Methods). (Center) Comparison of target selection performance (% correct) with PLTi intact (saline-teal) and PLTi silenced (CNO-red) in congruent and incongruent task conditions, n=6 PV-Cre mice; each line - one mouse. (2-way RM-ANOVA, p=0.00022 for Sal vs CNO followed by HBMC between Sal and CNO for % correct: Congruent - p=0.13, t(5)=1.8; Incongruent - p=0.002, t(5) = 5.6, paired two-tailed t-tests; (see Supplementary Table S1) (RT results in Figure 3.) (Right) Percentage correct (mean +/- s.e.m) for the congruent and incongruent conditions with weak flankers, saline vs CNO administration (2-way RM-ANOVA, p=8.52*10^-9^ for Sal vs CNO followed by HBMC between Sal and CNO for % correct: Congruent - p=0.228, t(10)=1.286; Incongruent - p=8.3*10^-4^, t(10) = 4.708, paired two-tailed t-tests; (see Supplementary Table S1). Box plots show the distribution of data along corresponding axes for comparison, demonstrating the large disruption in performance between saline and CNO for incongruent trials but not different in performance for congruent trials. **(d)** (Left) Abbreviated schematic of flanker task with task irrelevant flanker, and of bilateral chemogenetic virus injection in PLTi (Methods). (Right) Comparison of target selection performance (% correct) with weak task-irrelevant flankers with PLTi intact (saline) versus PLTi silenced (CNO), n=3 PV Cre mice (mean +/- s.e.m) (p=0.66, paired two-tailed t-test). Ghosted data is data from Figure 2c right for comparison purposes. See also Supplementary Figure 2h.

The flanker task is well established in humans^41-43^ for studying the guidance of behavior by information available at one spatial location at the expense of information available at all others. Specifically, it is known that flankers can outcompete the target in priority and draw attention away from it. In the incongruent condition, such attentional capture causes the flanker to guide behavior instead of the target, and results in systematic errors in performance (humans^41,42^; mice^14^). By contrast, in the congruent condition, such distraction by the flanker does not produce systematic errors in performance (% correct, nor reaction time) because the flanker contains the same information as the target^41,42,14^. Thus, any errors in the congruent condition are the result of nonspecific factors such as limits in learning stimulus feature discrimination, in reporting it correctly, and/or due to the selective processing of information elsewhere altogether. Comparison of performance in incongruent versus congruent trials for any given target and flanker priority, then, yields reliable metrics, specifically, of the extent to which selective spatial information processing, i.e., selective spatial attention, at the target location versus at the flanker location guides behavior.

In-line with these expectations^14,41,42^, we found that mouse target selection performance was lower in the incongruent flanker condition (pooled across flanker contrasts) compared to the congruent flanker condition (Figure 2b left and middle); performance was not significantly different between the congruent flanker and target-alone conditions (Supplementary Figure 2b). Notably, in just the trials in which the flanker contrast was *weak*, there was no difference in performance between incongruent and congruent conditions (Figure 2b right, % correct - gray filled circles; Supplementary Figure 2e, behavioral d-prime), consistent with behavior being predominantly guided by the high priority target in both cases, with errors reflecting non-specific factors. However, in trials in which the flanker contrast was *strong*, performance in the incongruent condition was significantly worse than in the congruent condition (Figure 2b right, % correct – black filled circles; behavioral d-prime, Supplementary Figure 2e; RT on error trials, Supplementary Figure 2e), consistent with the high priority incongruent flanker guiding behavior more frequently.

These findings were best explained by selective guidance of behavior by the incorrectly selected incongruent flanker because our results were able to rule out the two leading alternative explanations^14^. First, a purely low-level, motor-related explanation for the behavioral results was ruled out by the absence of head orienting differences between task conditions (Supplementary Figure 2c), and an absence of saccadic eye-movements during task-related epochs of trials (Supplementary Figure 2d). Second, an explanation of the flanker and target being perceived as a single composite stimulus, rather than two discrete, competing ones was ruled out by the lower accuracy in high contrast-incongruent trials compared to low contrast-incongruent trials (Figure 2b right, % correct; Supplementary Figure 2e, behavioral d-prime; see also^14^). Together, our results confirmed the validity of this touchscreen flanker task for studying selective spatial attention in freely moving mice.

We next tested the necessity of PLTi for this function. We bilaterally silenced the PV+ PLTi (“PLTi silencing”) in freely moving PV-Cre mice performing the flanker task using chemogenetics (Figure 2c left; administration of chemogenetic ligand CNO vs. saline to mice expressing the inhibitory chemogenetic receptor, hM4Di, selectively in PLTi; Methods). In the incongruent flanker condition, we found that PLTi silencing dramatically impaired target selection (trials pooled across flanker contrasts; Figure 2c middle, CNO administration, red data vs. saline administration, teal data); performance in congruent flanker trials was unaffected (Figure 2c middle, red vs. teal data). Notably, bilateral PLTi silencing caused a significant drop in performance even in trials with a weak flanker in the incongruent condition, but not the congruent condition (Figure 2c right, % correct; Supplementary Figure 2f, behavioral d-prime), consistent with increased distractibility and incorrect behavioral guidance by even weak incongruent flankers. The observed deficits were not due to non-specific effects of the chemogenetic ligand, CNO: administering CNO in mice expressing a reporter but not the chemogenetic receptor produced no discernible impact on performance (Supplementary Figure 2g; Methods). Thus, bilateral silencing of PLTi neurons disrupted spatially selective information processing for guiding behavior.

Considering that PLTi is a brainstem nucleus, positioned early in the neural processing hierarchy, we wondered whether these deficits were the result of PLTi neurons processing just bottom-up salience^1^, or whether it could be the result of PLTi neurons processing stimulus priority^42,43^ (top-down, goal-driven relevance + bottom-up, stimulus-driven salience). We reasoned that should PLTi neurons process just bottom-up salience, then silencing them in a task with an equally salient, but task-irrelevant flanker should also produce hyperdistractbility. To test this, we examined the effect of bilateral PLTi silencing in a subset of mice performing a modified version of the flanker task with just one change: the oriented flanker (distractor or ‘foil’) was replaced by a rectangular patch of equivalent luminance (task-irrelevant distractor; Figure 2d left; Methods). Strikingly, in the trials with weak flankers, we no longer found discernible deficits in performance in the task-irrelevant case in contrast with the task-relevant case (Figure 2d right vs Figure 2c right, %correct; Supplementary Figure 2h, behavioral d-prime; there was also no increase in RT following PLTi silencing - Supplementary Figure 2h). These results demonstrated that PLTi neurons are required for priority-dependent spatial target selection.

### PLTi is a dedicated brain module for attentional target selection

To what extent is the PLTi involved in attentional target selection for guiding behavior versus in other sensorimotor functions? This question is especially germane considering that the disruption of brain areas implicated thus far in selective spatial attention, such as fronto-parietal^6,9^ and midbrain^8,10^ areas, have been shown to also impair visual perception (i.e. perceptual sensitivity of single targets)^24-26^, motor plan selection (i.e., selection among orienting responses)^29-31^, and motor execution (i.e., trajectories and timing of orienting movements)^1,8-10,12^. Importantly, might potential impairments in any of these functions, rather than, specifically, in spatial target selection, explain the deficits observed in the flanker task following PLTi silencing? The bidirectional connectivity of PLTi neurons with the SC (Figure 1b and 1c), a major sensorimotor structure involved in all three of these other functions^8,44^ makes it critical to answer these questions.

First, we tested if PLTi neurons are involved in visual perception. Results from the target alone trials of the flanker task (same experiments as in Figure 2c) revealed that bilateral PLTi silencing did not significantly alter visual discrimination of single targets (Supplementary Figure 3a). To test if there may be a contrast-dependent role for PLTi on visual perception, we trained a subset of mice on a single target discrimination task in which the contrast of the target (oriented grating; same as in the flanker task) was varied systematically (Figure 3a; Methods). Here as well, we found that bilateral PLTi silencing in mice performing this task did not alter visual discrimination performance (Figure 3a middle) or perceptual sensitivity (Figure 3a right).

**Figure 3.**
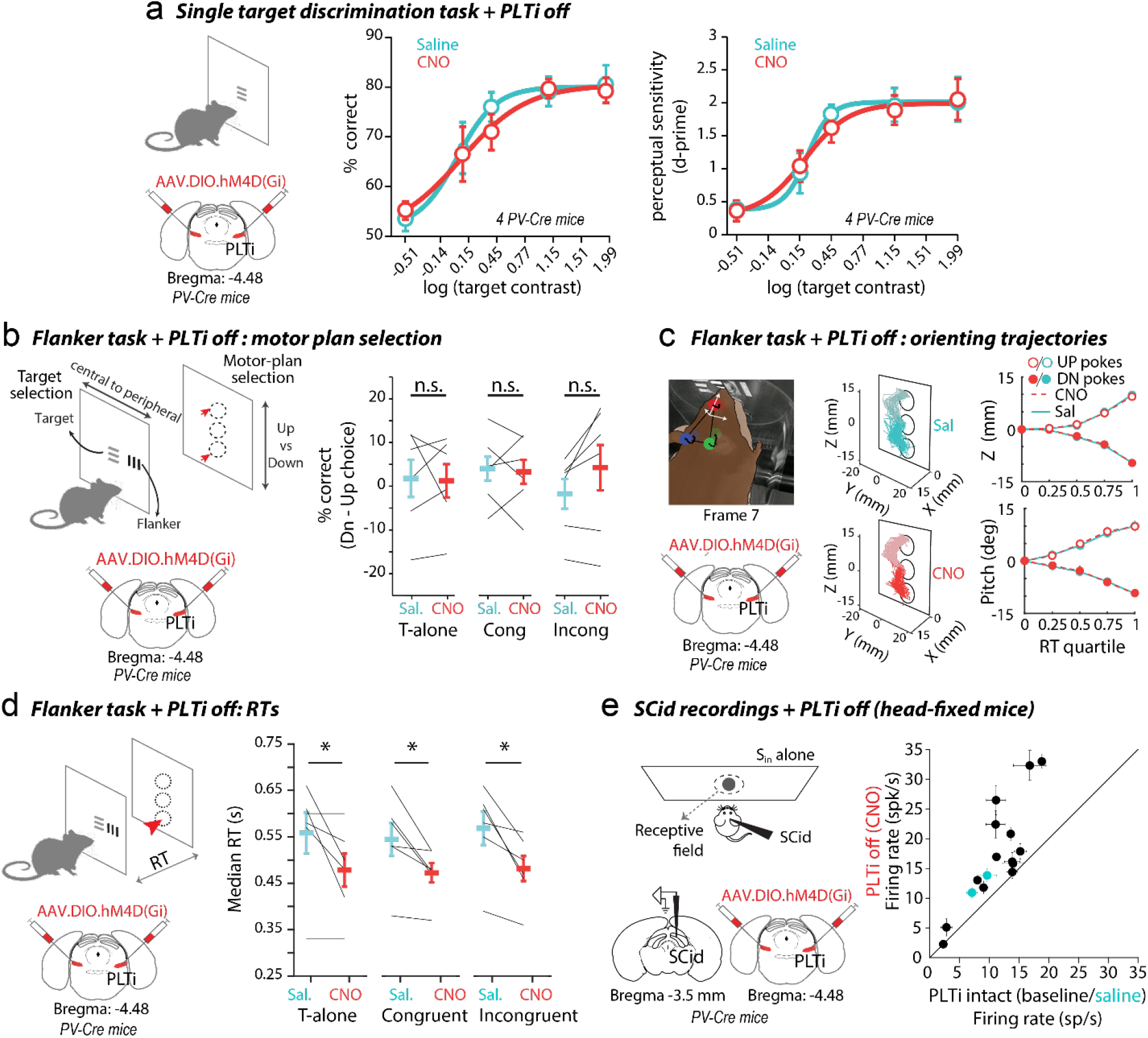
PLTi controls attentional target selection without affecting sensory perception, motor plan selection, or task-specific orienting movements. **(a)** (Left) Abbreviated schematic of single target discrimination task, and of bilateral chemogenetic virus injection in PLTi of PV-Cre mice (Methods). (Center and right) performance (% correct – center; p=0.81, two-way RM-anova for saline vs CNO, see Supplementary Table 1) and perceptual sensitivity (right; p=0.78, two-way RM-anova, see Supplementary Table S1) for the discrimination task as a function of contrast in saline (control, teal) and CNO (PLTi silenced, red) conditions, n=4 mice; mean +/- s.e.m. Solid lines – best sigmoidal fits (Methods). (b) (Left) Schematic of flanker task highlighting dissociation of target selection (left vs. right) and motor-plan selection (up vs. down) locations, and of bilateral chemogenetic virus injection in PLTi of PV-Cre mice (Methods). (Right) Difference in rate of selection of down vs. up choices in saline and CNO conditions. (target-only, p=0.39, t(5)=16.12, two-tailed t-test; congruent, p=0.93, t(5)=6.49, two-tailed t-test; incongruent, p=0.2, t(5)=13.92, two-tailed t-test, followed by HBMC for multiple comparisons in each case; n = 6 PV-Cre mice; same experiments as in Figure 2c. Conventions as in Figure 2b. (c) (Left top) Snapshot of mouse orienting movements in a nose-touch up-choice trial, for frame #7 in a trial. Red, blue and green filled circles – head-tracking labels generated by automated deep learning tool (Methods). Black dotted lines – movement trajectories across all tracked frames. White arrows – egocentric coordinate axis constructed using the three labeled markers; see also Supplementary Figure 3b. (Left bottom) Schematic of virus injection in PLTi of PV-Cre mice. (Center) Reconstructed head movement trajectories in all saline (top, cyan) and all CNO (bottom, red) trials for an example session. Lighter colors – trials in which mouse nose-touched upper port. (Right) Head position of mouse tracked over the course of each trial; plotted as mean +/- s.e.m across trials, with each trial split into RT quartiles; (Top) Z-coordinate; (Bottom) pitch angle. Cyan-saline, red-CNO, open (closed) circles – trials with nose-touched to upper (lower) port. See Supplementary Figure 3b for X, Y coordinates and roll, yaw angles. P>0.2, for each X, Y and Z coordinates; p>0.3, for each roll, yaw and pitch angles; two-way RM-anova in each case, see Supplementary Table S1. (d) (Left) Abbreviated schematic of flanker task, and of bilateral chemogenetic virus injection in PLTi of PV-Cre mice. (Right) Median reaction times across flanker task conditions; saline (teal) vs. CNO (red); conventions as in Figure 2c. (left, target-only, p=0.04, t(5)=2.81; middle, congruent, p=0.024, t(5)=3.11; right, incongruent, p=0.01, t(5)=3.51 – two-tailed t-tests in each case followed by HBMC). (e) (Left top) Schematic of experimental setup for extracellular recordings in SCid of passive, head-fixed mouse. Sin: single stimulus presented inside receptive field of SCid unit. (Left bottom) Schematic of recordings in SCid and chemogenetic virus injection in PLTi of PV-Cre mice. (Right) Mean evoked firing rates (+/- s.e.m) of SCid neurons in baseline/saline (black/teal) vs. CNO (red) conditions; n=14 units, p=0.001, t(15)=9.418, paired two-tailed t-test).

Second, we tested if PLTi neurons are involved in motor plan selection. In our flanker task, animals selected between motor plans for responding to the upper port versus the lower port. Therefore, we compared relative response choice rates to the upper versus lower response ports. This comparison served as a reliable test for motor plan selection because our task design dissociated the locations of behavioral response from the locations of sensory stimulus presentation (Figure 3b left). We found that bilateral PLTi silencing had no effect on upper versus lower choice rates in any condition of the flanker task (Figure 3b right). Third, we tested if PLTi neurons are involved in the execution of orienting movements, namely, their trajectories and their timing. We first tracked the 3-dimensional head-orienting trajectories of mice performing the flanker task using calibrated video cameras (Figure 3c left and center; same experiments as in Figure 2c; Methods). We found that bilateral silencing of PLTi neurons did not produce any detectable changes in movement trajectories during touchscreen interactions: either in 3D head displacement (Figure 3c right top and Supplementary Figure 3b), or in 3D head orientation (Figure 3c right bottom and Supplementary Figure 3b).

To test PLTi’s involvement in controlling the timing of orienting movements, we analyzed mouse reaction times (RTs) in the flanker task (same experiments as in Figure 2c). We found that PLTi inactivation caused *faster* (decrease in) median RTs, and did so in *all three* task conditions (T-alone, congruent and incongruent; Figure 3d). We wondered if this might present an alternate explanation (to disruption in target selection) for our results on response accuracy following PLTi silencing (Figure 2c). This is because, it is established that faster RTs accompanied by lower accuracies can be well-explained simply through the speed-accuracy tradeoff. If true, a change in the operating point along the speed-accuracy tradeoff that produced faster RTs would necessarily predict lower accuracy in *all* task conditions. However, following PLTi silencing, we found that accuracy was lower only in the incongruent condition but was unaffected in the congruent and single target condition (Figure 2c and Supplementary Figure 3a). This ruled out a change in speed-accuracy tradeoff and the corresponding confounding explanation.

How, then, might bilateral PLTi silencing cause speeding up of RTs across task conditions? To gain insight into this question, we turned to drift diffusion modeling (DDM^45,46^) of RT distributions from the flanker task (Methods, Supplementary Figure 3c). We found that PLTi silencing caused a reduction of the threshold parameter of the DDM (threshold ‘a’; Methods; Supplementary Figure 3d-f) in *each* of the flanker task conditions following PLTi silencing, revealing a non-specific decrease in movement initiation threshold. Mechanistically, how might PLTi silencing affect movement initiation thresholds? We hypothesized that such a decrease in movement thresholds would be supported by a non-specific increase in evoked firing rates of SCid neurons following PLTi silencing. This is because SCid, which receives tonic inhibition from PLTi (Figure 1), is well-known to causally control species-relevant orienting movements across vertebrates^44,47^. Specifically, firing rates of SCid neurons are known to be related to thresholds for movement initiation^48,49^. To test this hypothesis, we performed electrophysiological recordings in SCid of passive (awake and head-fixed) mice (Figure 3e left, Methods). We found that bilateral PLTi silencing caused an increase in evoked firing rates of SCid neurons to single stimuli even in passive mice (Figure 3e right, Methods). Thus, the blanket increase in SCid firing rates due to PLTi silencing-induced disinhibition of SCid accounted for the non-specific speeding up of reaction times across task conditions.

Together, these results yielded two powerful findings. First, that PLTi controls spatial target selection *purely* by controlling competitive interactions between the target and distractor (because PLTi silencing leaves intact single-target visual perception - Figure 2c and 3a)). Second, that PLTi is a, *dedicated* module for spatial target selection amidst distracters (because PLTi silencing leaves intact visual perception, motor plan selection, and task-specific timing of orienting movements, unlike other areas implicated thus far in selective spatial attention^9,10,12^).

### PLTi controls the cognitive decision boundary separating the attentional target from distractor

Thus far, our analyses have examined the effect of PLTi silencing on behavior in the flanker task at extreme values of relative priority (very weak flanker or very strong flanker; Figure 2b). While this approach has revealed the necessity of PLTi for target selection among distracters, it leaves unaddressed our understanding of target selection processes at intermediate values of relative priority at which discrimination between the target and distracters is typically the hardest. Indeed, the question of how neural circuits control selection of a target among distracters across the full range of relative priorities - a question that is particularly germane to behavior in dynamic natural environments - has remained largely unstudied; prior work has typically assessed performance only at one or a few values of relative priority^21,27,50^. As a result, the mechanistic implementation of the theorized winner-take-all^51,52^ target selection is unknown^53^. Our flanker task, with its systematic parameterization of flanker priority, is well-suited to study this question.

As a first step, we analyzed mouse performance in the flanker task as a function of flanker priority when PLTi was intact (Figure 4a left and Figure 2a; “baseline” dataset from Figure 2b). Performance in congruent flanker trials was largely unaffected by flanker contrast (relative priority; Figure 4b, gray points – example animal; Supplementary Figure 4b – all animals; results similar to humans^41,42^ and mice^14^). However, in the incongruent flanker trials, target selection accuracy (% correct) decreased as a function of flanker contrast, and this psychometric selection curve was well-fit by a sigmoid^14^ (Figure 4b, black points and curve; Supplementary Figure 4a,b; Methods).

**Figure 4.**
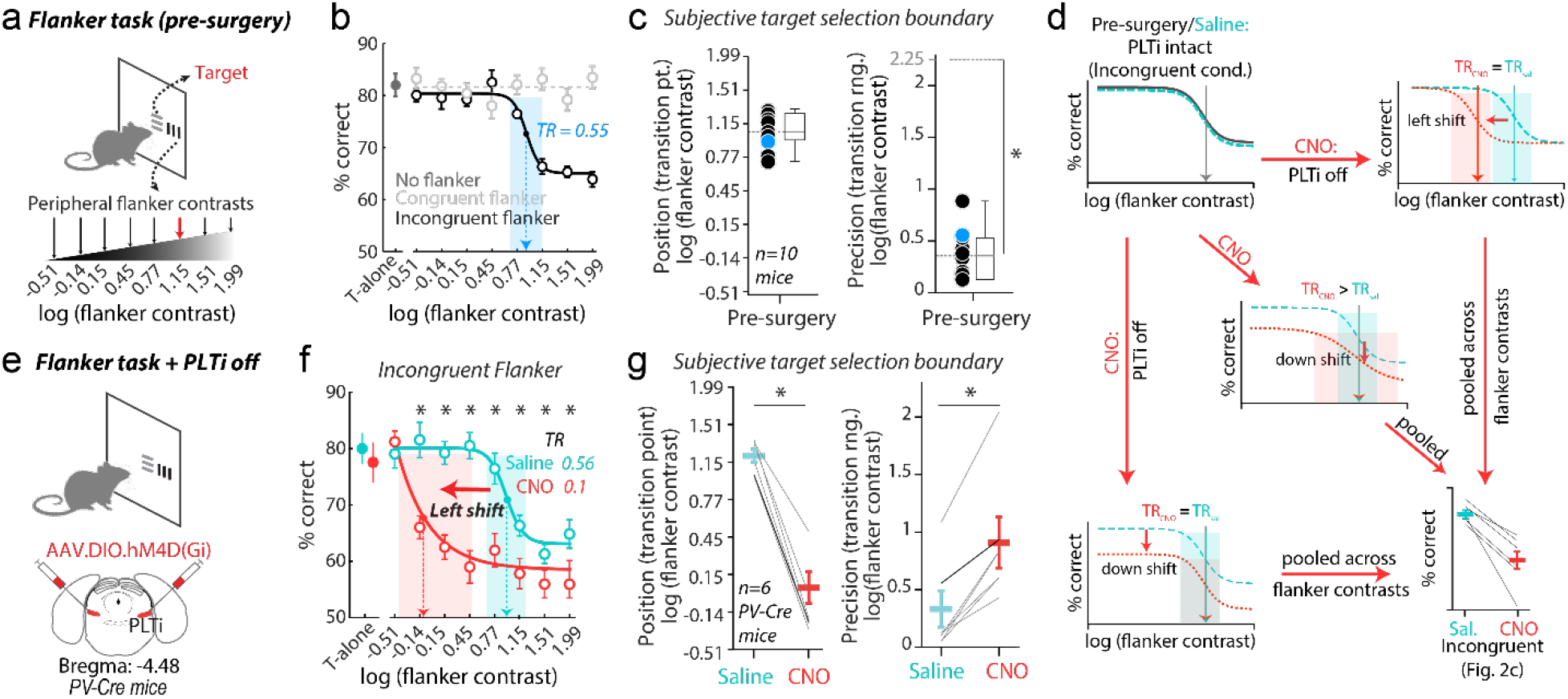
PLTi controls the cognitive decision boundary between attentional target and distractor. **(a-c)** Position and precision of behavioral target selection boundary in flanker task (incongruent flanker trials); data from same experiments as in Figure 2b. **(a)** Abbreviated schematic of flanker task showing flanker contrasts (arrows) along increasing (log) scale. **(b)** Target selection performance (% correct) of example mouse as a function of flanker contrasts, data from incongruent (black open circles), congruent (gray open circles), and target-alone (gray solid circle) conditions; mean +/-s.e.m. Dashed light gray line: mean of congruent data. Black solid line: best sigmoidal fit to incongruent data (Methods). Blue arrow and black solid circle: position of decision boundary (TP-transition point; Methods). Blue shading: precision of decision boundary, quantified by TR (transition range). **(c)** Summary plots characterizing the (left) position of the decision boundary (TP of the response profile), and (right) precision of the decision boundary (inversely related to TR); p=1.57*10^-9^, t(9)=24.37, paired two-tailed t-test; 2.25 = transition range of the least precise, linear decision boundary; n=10 mice. Each circle represents data from one mouse; blue circle: from example mouse in panel b. **(d-g)** Effect of bilateral PLTi silencing on position and precision of decision boundary in flanker task (incongruent flanker trials); data from same experiments as in Figure 2c. **(d)** (schematics) (Top-Left): baseline and saline target selection performance curves for incongruent trials as a function of flanker contrast. (Top-Right, Center and Bottom-Left): hypothetical target selection performance curves after bilateral PLTi silencing, showing potential changes in the position and/or precision of the decision boundary, each of which, when pooled across flanker contrasts, could explain our results in Figure 2c. (Bottom-Right) Panel from Figure 2c. **(e)** (Top) Abbreviated schematic of flanker task, and (bottom) of bilateral injection of inhibitory chemogenetic virus in PLTi of PV-Cre mouse. **(f)** Target selection performance (% correct) of example PV-Cre mouse as a function of flanker contrasts in incongruent trials; saline (control, cyan) vs CNO (PLTi silencing, red) conditions, black asterisks: p<0.05, RM-anova and post-hoc t-tests with HBMC; each point is mean +/-s.e.m; other conventions as in panel b. CNO disrupts position (leftward shift – red arrow) and precision (lower TR) of decision boundary. **(g)** Summary plots showing effect of PLTi silencing on (right) position of the decision boundary (TP); p=6.04*10^-7^, t(5)=4.78, paired two-tailed t-test), and (right) precision of the decision boundary (inversely related to TR); p=0.006, t(5)=4.55, two-tailed t-test. n=6 PV-Cre mice; saline (control): teal; CNO (PLTi silencing): red.

The mid-point of this sigmoid, defined as the flanker contrast at which accuracy was half-way between the maximum and minimum (‘transition point - TP’; Figure 4b – example animal, Supplementary Figure 4a – all animals; dots and arrows), signaled a qualitative transition in the extent to which the target stimulus was favored for selection and behavioral guidance: for lower values of flanker contrast, behavior was guided almost exclusively by the target, whereas for higher values of flanker contrast, the flanker outcompeted the priority of the target stimulus and guided behavior significantly more often^14,42,54,55^. This transition point, therefore, signaled the subjective *cognitive decision boundary* – the behavioral threshold along the axis of relative priority that separates the highest priority target from lower priority distracters for each animal, thereby dictating target selection (Figure 4b, black solid dot; Supplementary Figure 4a). Across animals, this subjective decision boundary occurred on average when flanker contrast was 1.1 (units of log(contrast), s.e.m = 0.1 units; Figure 4c left).

Separately, the steepness of this sigmoid, quantified by the range of contrast values over which the responses dropped from high to low levels of accuracy (‘transition range - TR’; Figure 4b, shading – example animal, Supplementary Figure 4b, shading –all animals; Supplementary Figure 4a – schematic; Methods;^56^), signaled the *precision* of the subjective decision boundary: narrower transition ranges represented a more categorical selection of the target among distracters (schematic Supplementary Figure 4a;^56,57^). Across animals, the transition ranges were narrow, and significantly smaller than that of a linear psychometric curve (Figure 4c right; 0.37 +/- 0.07 log(contrast); Supplementary Figure 4a, black line; Methods). Thus, the decision boundary in our flanker task was categorical (winner-take-all-like and highly precise;^53,58^). These results established that the flanker task provides a succinct, quantitative description of target selection amidst distracters across the full range of relative priorities. It allows quantitative measurement of the cognitive decision boundary through its two defining computational attributes - position and precision along the axis of relative priority.

With this foundation, we next asked whether the PLTi is involved in the control of either the position or the precision of the cognitive decision boundary. The answer to this question cannot be predicted from the effects reported thus far using pooled contrasts (Figure 2c) because they can arise from multiple distinct combinations of effects on position and precision including even, no effect on either (Figure 4d, schematic). We, therefore, examined the effects of PLTi silencing on the relative priority-dependent psychometric selection curves (Figure 4e). We found that in the incongruent flanker trials, PLTi silencing caused both the position and the categorical precision of the decision boundary to be impaired severely (Figure 4f – example animal; Supplementary Figure 4c – summary across animals). The subjective decision boundary was shifted leftward (Figure 4f, red arrow; Figure 4g left and Supplementary Figure 4c), indicating that lower priority flankers incorrectly outcompeted the target and guided behavior. Separately, the decision boundary was also less precise, i.e., the response profiles became less categorical (Figure 4f, red vs teal shaded area; Figure 4g right and Supplementary Figure 4c). Performance in the congruent trials was unaffected by bilateral PLTi silencing (Supplementary Figure 4c). These results were not due to non-specific effects of the chemogenetic ligand (CNO): administering it in mice not expressing the chemogenetic receptor produced no discernible effects in the incongruent flanker condition (Supplementary Figure 4d-f). Thus, PLTi controls the cognitive decision boundary along the axis of relative priority, specifying both its subjective position and categorical precision.

### PLTi controls the categorical neural selection boundary between competing stimuli in the SC

How, mechanistically, does the PLTi exert control over behavior, and specifically, on the position and categorical precision of the cognitive decision boundary? We hypothesized that PLTi may do so by shaping competitive interactions and selection signals in the SCid^21^ (Figure 1). To test our hypothesis, we extended our recordings in head-fixed, passive mice (Figure 5a top and bottom; Figure 3e) to examine the nature of competitive interactions in mammalian SCid. We measured responses of SCid neurons to a stimulus competition protocol inspired by our flanker task (and identical to the protocol used in previous avian work^56^; Figure 5a top, Methods). Briefly, we presented one stimulus, S_in_, a dot of fixed strength (loom speed), within the spatial receptive field (RF) of the SCid neuron (typically in the binocular visual field), and a second, competing stimulus, S_out_, of varying strength far outside the RF (typically in the peripheral visual field; Figure 5a top, Methods)^56^. We found that responses of SCid neurons to S_in_ decreased in a sigmoidal manner with increasing salience of competitor S_out_ (Figure 5b, example neuron). The transition point (TP) of the sigmoid (i.e., position of the *neural* selection boundary in SCid for signaling the most salient stimulus) occurred at a relative salience of zero between the competing stimuli (Figure 5b right, example neuron – black dashed arrow; Figure 5c left, summary across neurons). In parallel, the transition range (TR) of the response profile (signaling precision of the *neural* selection boundary in SCid for signaling the most salient stimulus) was narrow (Figure 5b right, example neuron – shaded region; transition range = 0.7 deg/s; Figure 5c right, summary across neurons). Thus, the neural selection boundary separating the most salient stimulus from the others is encoded explicitly by SCid neurons in an accurate and precise (winner-take-all-like) manner (matching previous reports in birds;^56,59^).

**Figure 5.**
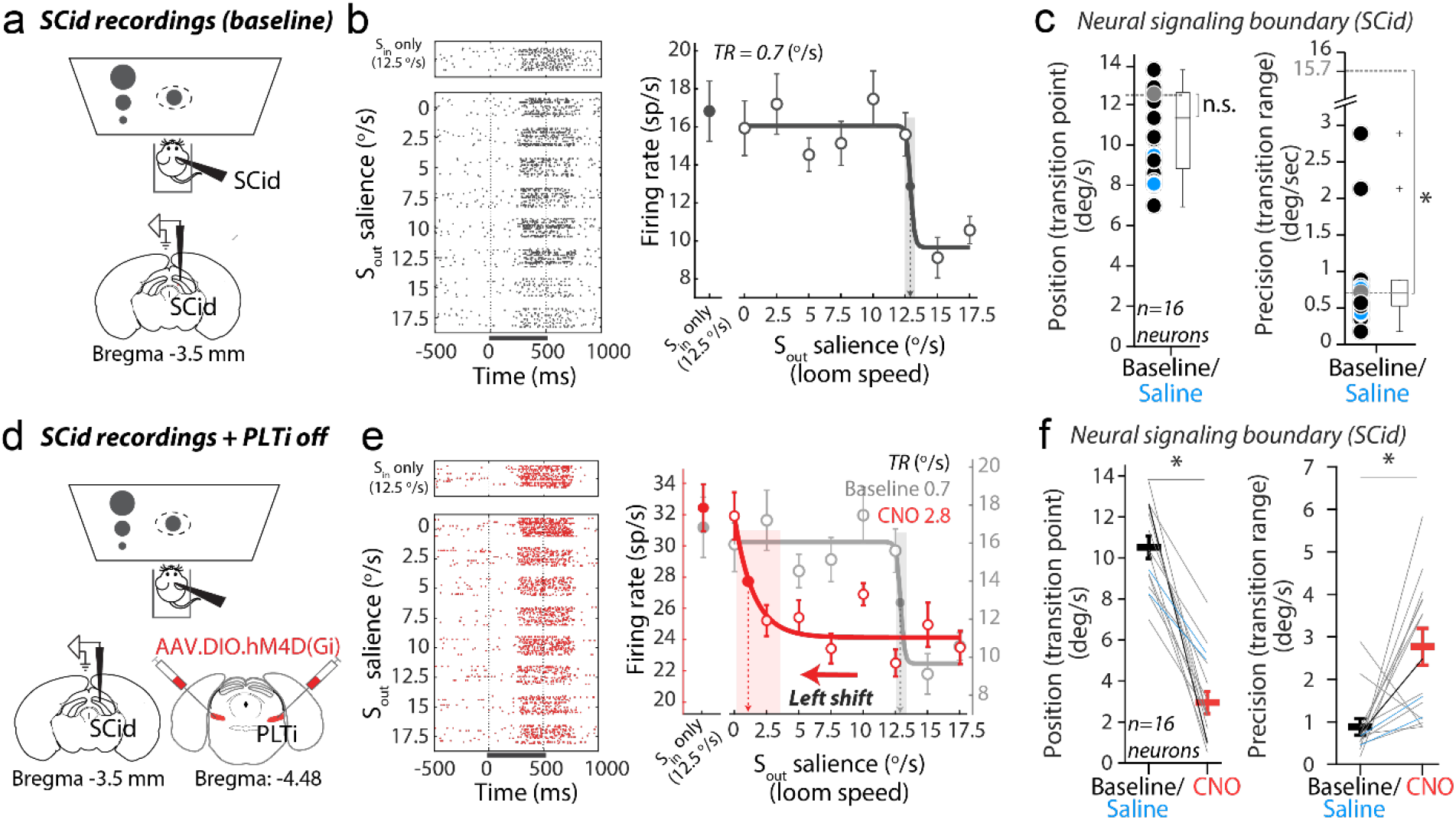
PLTi controls the neural selection boundary between competing stimuli in the SC. **(a-c)** Position and precision of neural boundary in SCid explicitly signaling the most salient stimulus. **(a)** (Top) Schematic of stimulus protocol for characterizing competitive interactions in SCid using two stimuli: Sin – receptive field stimulus and Sout – competing peripheral stimulus (Methods). (Bottom) Schematic showing location of extracellular recordings in SCid (Methods). **(b)** Responses of example SCid unit to competition protocol. (Left) Raster plot, gray bar below raster: stimulus duration. (Right) Firing rates (mean +/- s.e.m). Open circles: responses to Sin and Sout, solid line: best sigmoidal fit; solid circle: responses to Sin alone. Arrow: position of neural signaling boundary, close to relative salience=0, which is when Sin and Sout are equal in strength at 12.5 (deg/sec); shading: precision of neural signaling boundary. **(c)** Summary plots showing transition points of neural selection boundary (position; (left); p=0.125, t(15)=8.87, two-tailed t-test against relative salience of 0, which corresponds to 12.5 deg/sec indicated by the gray dotted line; Methods) and transition ranges of neural signaling boundary (precision; (right); p=1.33*10^-18^, t(15)=75.9, two-tailed t-test against the TR – 15.7 deg/sec = transition range of the least precise, linear response profile; n=16 units; Methods). **(d-f)** Effect of bilateral PLTi silencing on position and precision of neural signaling boundary in SCid; panels and conventions as in a-c. **(d)** (Top) Schematic of stimulus protocol (same as in panel a). (Bottom) Bilateral injection of inhibitory chemogenetic virus in PLTi of PV-Cre mouse. **(e)** Responses of same example SCid unit as in panel b to competition protocol, but with PLTi silenced (CNO, red). (Left) Raster plot. (Right) CNO disrupts position (leftward shift, arrows) and precision (lower TR, shading) of neural signaling boundary, wider shading (pink) implies lower precision or less categorical. **(f)** Summary plots of transition points (position; (left), p=1.58*10^-7^, t(15)=5.95, paired two-tailed t-test; Methods) and transition ranges (precision; (right), p=0.005, t(15)=3.34, paired two-tailed t-test) of neural signaling boundary with PLTi silenced; n=16 units (lines; Methods).

We next examined the effect of bilateral PLTi silencing on competitive interactions in the SCid (Figure 5d). Strikingly, PLTi silencing disrupted both the position and categorical precision of the *neural* selection boundary (Figure 5e). It caused a leftward shift of SCid response profiles (shift in position-TP), indicating that a weaker S_out_ was sufficient to outcompete and powerfully suppress responses to S_in_ (Figure 5e right, example neuron; Figure 5f left, summary across neurons). PLTi silencing also caused the response transitions to become substantially wider (less categorical; Figure 5e right example neuron; Figure 5f right, summary across neurons), thus decreasing the precision of the neural selection boundary. (Consistent with Figure 3e, we also found an overall increase in stimulus-evoked SCid firing rates, Supplementary Figure 5a). These effects of PLTi silencing on the SCid’s *neural* selection boundary were identical to the effects of PLTi silencing on the *cognitive* decision boundary (Figure 4f,g). Considering the established necessity of intact competitive stimulus interactions in the SCid for target selection^21,30^, these results reveal that PLTi’s shaping of competitive interactions in SCid is a major mechanistic path by which PLTi sets the position and categorical precision of the decision boundary and drives selective spatial attention.

## Discussion

In this study we have discovered that an ancient^32,37,38,60,61^ group of PV+ inhibitory neurons in the mouse brainstem, PLTi, controls how mice sift through the barrage of information across space to identify the most important piece for guiding behavior.

### Behavioral task designs in (head-fixed) primates versus in (freely behaving) mice for studying selective spatial attention

Overt changes in gaze direction do not confound our interpretations. Shifts in the spatial locus of selective visual information processing can occur overtly through changes in gaze direction (head + eye-in-head movements), or covertly. Studies of selective spatial attention in (head-fixed) primates typically adopt an experimental design that explicitly minimizes (or penalizes) such overt shifts in gaze. This has allowed for unambiguous mapping of measured neural correlates or causal mechanisms for target selection onto higher-level cognitive processes over lower-level motor factors related to overt gaze shifts^12,21,26^. Our study in freely moving mice does not explicitly penalize overt shifts in gaze. Notably, however, unlike in primates, it has been shown in mice, and specifically in freely behaving mice, that eye movements in the head during goal-directed behavior sub serve primarily convergence and compensation, rather than saccadic acquisition of targets^62^. Our results (Supplementary Figure 2) directly confirm this finding during performance in our flanker task, demonstrating the absence of significant saccadic eye movements during task epochs. Moreover, our results demonstrate not only the absence of significant differences in head movement trajectories between task conditions (Supplementary Figure 2), but also the absence of changes in head movement trajectories upon PLTi silencing (Figure 3c and Supplementary Figure 3). Taken together, these results in freely behaving mice support that changes in overt gaze shifts do not underlie PLTi-silencing induced changes in behavior in our flanker task, aligning our results to the interpretational framework of classical monkey experiments.

### An evolutionarily conserved nucleus for selective spatial attention

Attentional target selection requires identification of the spatial location with the highest priority information, rather than with just the most physically salient information. Our results demonstrate that PLTi integrates top-down (behaviorally relevant) inputs as well as bottom-up (physically salient) inputs to selectively guide behavior (Figure 2). They provide direct evidence that attentional target selection is achieved by brain networks implementing stimulus competition based on the combined salience and relevance of stimuli, rather than only by separate (and potentially competing) networks implementing only salience-based (bottom-up) or relevance-based (top-down) selection^63,64^. PLTi’s necessity for spatially selective information processing to guide behavior, despite its evolutionarily antecedence and its early station in the neural processing hierarchy^32,33,65^, is also striking. They promote a re-examination of traditional notions of an exclusively fronto-parietal neural basis of this crucial executive function. Moreover, they position the relatively unknown group of inhibitory neurons as the missing link for understanding the conserved neural basis for attentional target selection across vertebrate taxa^13,20^.

### Explicit instantiation of cognitive decision boundary

The computational roles of PLTi sub-serving target selection are sophisticated. The absence of an effect of PLTi silencing on the perception of single targets (Figure 2c and 3a) reveals that PLTi controls target selection via distractor suppression, specifically by orchestrating competitive interactions between the target and distracter(s). Our results reveal key details about the specific computations related to competitive stimulus interactions that PLTi regulates. First, they reveal that PLTi controls the *position* of the decision boundary, i.e., the point of subjective equality of the priorities of the target and distractor (Figure 4). Notably, the observed shift of this boundary disfavoring the target and affording undue competitive advantage to the distractor, i.e. hyperdistractibility, parallels attentional phenotypes in neurodivergent conditions (including ADHD and schizophrenia;^4,5^). Second, our results show that PLTi also controls the degree to which selection of the next spatial target for behavior is winner-take-all-like (as a function of relative stimulus priority) (Figure 4). In the natural world replete with sensory ambiguity and response variability, such categorical psychometric response profiles boost reliability in selecting the highest priority target^53,66^ - an ethologically adaptive feature. Consequently, these results offer, for the first time to our knowledge^6,8-12^, a neurobiological circuit mechanism for the explicit control of the position and precision of the decision boundary, and of winner-take-all-like selection, in decision-making tasks. (We note that the attributes of the decision boundary described above are conceptually distinct from the frequently studied “criterion” and “perceptual sensitivity”^50^: while the former attributes require parameterized investigation of selection across a range of relative-priority values, the latter characterize discrimination performance at each value of relative priority, and can vary with it).

### Control of the SC relative priority map to drive behavior

The powerful computational contributions of PLTi to the control of selective spatial attention raise the question of the mechanistic path(s) of action that realize them. Our results demonstrate that PLTi, which functionally inhibits the midbrain SC via long-range projections (Figure 1d), controls competitive interactions among stimulus representations in the SCid (Figure 5). On the one hand, this resolves the long-standing mystery regarding the *source* of inhibition for competitive interactions in the mammalian SCid^67-69^, a major player in sensory-guided behavior, spatial decision-making and selective spatial attention^8,10^, and in turn, discovers a mechanism for the formation of the map of relative priority in the SCid. On the other, the striking correspondence between the effects of PLTi silencing on the position and precision of the *neural* selection boundary signaled by SCid neurons, and the effects on the position and precision of the *behavioral* decision boundary expressed by the animal, indicates that PLTi’s shaping of responses of SCid neurons represents a major mechanistic path of its action on behavior.

### Specialized modules versus distributed implementations

Spatial target selection for selective attention is closely linked to the functions of sensory perception, motor plan selection and orienting behavior. Indeed, brain regions implicated in the control of spatial target selection, such as the fronto-parietal cortices^9,12^ and the midbrain superior colliculus^8,10^, play important roles also in these sensorimotor functions. Not surprisingly then, lesions or inactivation of these regions are accompanied by deficits in perceptual sensitivity to single targets, motor planning or gaze orienting^9,10,12,26^. In this study, however, we found that bilateral silencing of PLTi caused dramatic deficits in competitive target selection (Figure 2c) without producing discernible deficits in single target perception (Figure 3a), motor planning (Figure 3b) and task-related head-orienting movements (Figure 3c). These results reveal PLTi as a special, dedicated brain module for winner-take-all-like spatial target selection (over other related sensorimotor functions) during goal-driven behavior. Consequently, they support a structured, computationally driven organization of brain systems for higher cognitive functions^3,53,58^, in departure from recent suggestions of extraordinarily distributed neural substrates^70,71^.

### Open questions

Whereas our study has demonstrated PLTi’s necessity for target selection underlying selective spatial attention, the detailed mechanistic (structural and functional) logic of operation of the mammalian PLTi-SC circuit is unknown. Additionally, whether well-structured circuit modules similar to the PLTi exist within the fronto-parietal networks that are also known to be crucial for such spatial target selection^9,12^ is an intriguing open question. Finally, the functional logic by which brainstem (Figure 2,4 and 5;^20,72-75^), thalamic^11^ and fronto-parietal^6,9,12^ circuits work together to drive spatial target selection amongst distractors, remain to be understood. Addressing these questions will be essential for a mechanistic understanding not only of spatially selective information processing for behavioral guidance in healthy states, but also in disrupted states found in neuroatypical conditions.

## Acknowledgments

We thank Sarah Correas, Samrawit Getachew, Isabel Rios Pulgar, Sarah Saccal, Marvin Lee, Huiyao Chen, Elisa Herrera, Spencer Loggia, and Ronald Salazar for help with behavioral training of the mice, Spencer Loggia for help with the cameras, and Rohit Singh for help with eye-tracking. We thank Dr. Eric Garr for sharing protocols for CNO use. We thank James Garmon for engineering support. We are grateful to Dr. Howard Egeth, Dr. James Knierim and Dr. Kishore Kuchibhotla for helpful feedback on the manuscript.

## Funding

This work was supported by NIH F32EY032776 (NBK), NIH R34NS111653 (SPM), NSF 2047298 (CAREER award, SPM) and start-up funds from the Johns Hopkins University (SPM).

## Author contributions

SPM and NBK conceived the project and designed the experiments; NBK performed and supervised behavioral experiments and data analyses; NBK and QZ performed the electrophysiological experiments and analyses; AB, WKY and NBK performed anatomical experiments and analyses; SPM and NBK wrote the paper.

## Declaration of interests

The authors declare no competing interests.

## Data and materials availability

Code, data and materials used in the analysis will be made available upon reasonable request to the corresponding author.

## Notes

### Competing Interest Statement

The authors have declared no competing interest.

### Summary of Updates

Figure 2, Figure 4 and Figure 5 revised. Eye movement data using a novel non-invasive technique added.

